# Molecular Basis of Sperm Methylome Response to Aging and Stress

**DOI:** 10.1101/2024.11.12.623255

**Authors:** Olatunbosun Arowolo, Jiahui Zhu, Karolina Nowak, J Richard Pilsner, Alexander Suvorov

## Abstract

Changes in the sperm epigenome induced by age and/or stressors often follow common unexplained patterns affecting genes responsible for embryonic development and neurodevelopment. The stochastic epigenetic variation (SEV) hypothesis proposes that in response to stressors naturally variable methylation regions (VMRs) associated with morphogenic genes increase in methylation variation to diversify phenotypes and improve chances of survival of the genetic lineage. Here, we test predictions from the SEV hypothesis using mouse and rat sperm DNA methylation and other -omics data. We demonstrate that the context of DNA regions determines the response of sperm methylome to various factors rather than the stressors and/or timing of these factors. We propose a model explaining age/stress-dependent shifts in methylation in VMRs by an asymmetric increase in methylation variation of these regions. Because methylation variation in VMRs increases with age, sperm methylome response to stressors may be characterized as an acceleration of epigenetic aging.

## INTRODUCTION

Research conducted over the last two decades demonstrates that sperm DNA methylation is sensitive to reprogramming by a broad range of factors including in-utero undernourishment ^1^, paternal diet ^2^, childhood abuse ^3^, psychological stress ^4^, metabolic status ^5,6^, heat stress ^7^, environmental exposures during perinatal development or the preconception period ^8–13^, and aging ^8,9,14,15^ among others. After fertilization, parental-specific epigenetic marks of gametes undergo reprogramming to establish totipotency in the developing embryo. However, imprinted loci and certain classes of repetitive sequences escape this reprogramming event and contribute to the non-Mendelian form of inheritance ^16^. Sperm DNA methylation in other genomic loci may also resist reprogramming as reported in human ^17,18^ and rodent ^4,15,19,20^ studies. This evidence demonstrates that the final sperm methylome acquired during spermatogenesis is a significant channel for the transfer of inheritable information to the next generation ^21^.

Final DNA methylation profiles of spermatozoa are a result of epigenetic events that occur during spermatogenesis ^22^, including meiotic divisions of spermatogonia and spermatocytes associated with passive loss of methylation and *de novo* methylation ^23^. Reduction in global methylation of around 12-13% occurs in preleptotene spermatocytes and methylation is gradually reestablished during leptotene-pachytene stages so that the final DNA methylation patterns are obtained by the end of pachytene spermatocyte stage ^23–25^. A recent rat study identified the transition from elongating spermatids to late spermatids as another stage associated with changes in methylation of more than 5,000 DNA regions ^26^.

Published research demonstrates that changes in sperm epigenome mediate the effects of paternal factors on adverse outcomes in offspring. Some examples include the effects of advanced paternal age on life span reduction ^19^, altered vocal communication ^27^, reduced exploratory and startle behavior ^15^ in mouse offspring, and reduced fertilization rates in humans ^28^; metabolic dysfunction in mice induced by paternal psychological stress ^4^, paternal diet ^2^, paternal prediabetes ^5^, and paternal in utero undernourishment ^1^, and increased risk-taking behavior induced by paternal psychological stress in mouse offspring ^4^. Additionally, the role of sperm epigenetics in offspring health is suggested by population studies linking advanced paternal age (irrespective of maternal age) with poor pregnancy outcomes ^29^, lower odds of live birth ^30^, and adverse health of the offspring at later life ^31–33^, including increased susceptibility to the early development of cancer ^34^; and neurodevelopmental disorders such as schizophrenia ^35^ and autism ^32,33,36,37^. These changes in the sperm epigenome induced by age or various stressors are often explained by the accumulation of random epigenetic errors or epimutations ^38–41^. However, as more data on sperm epigenetics are becoming available, common patterns of epigenetic change are emerging and challenge the random nature of sperm epigenetic changes ^42^.

First, according to recent studies two epigenetic mechanisms, DNA methylation, and profiles of small non-coding RNA (sncRNA) undergo concordant changes in sperm. For example, Xie et al ^19^ demonstrated that in mice both age-dependent changes in DNA methylation and sncRNA in sperm were associated with major developmental and aging pathways. In our experiments with rats, age-dependent changes in sncRNA ^9^ and DNA methylation ^8^ were enriched in the same set of developmental pathways, including those identified by Xie et al. In another study, increased adiposity and reduced glucose tolerance in mouse offspring were equally induced by sperm and seminal plasma of fathers on low protein diet ^2^, suggesting that sncRNA cargo of extracellular vesicles in seminal plasma ^43–45^ and spermatozoa-specific epigenetic mechanisms transfer similar epigenetic information to the next generation.

Second, in a broad range of studies of the effects of various factors on sperm DNA methylation regardless of the factor analyzed (e.g., age, diet, chemical exposure, heat stress, psychological stress) and biological species (human, rat, mouse) genes associated with DMRs in sperm are almost always enriched for the same set of biological categories, including embryonic development, neurodevelopment, behavior, and metabolism ^1,3,5–8,10,11,14,18–20,46–49^. These conserved patterns of sperm epigenetic changes suggest that a purely stochastic error model cannot explain the shared changes in sperm epigenome and strongly suggests an evolutionary basis of this observed phenomena ^50^.

According to the evolutionary stochastic epigenetic variation (SEV) hypothesis, stress may increase variation in naturally variable methylation regions (VMRs) associated with development and morphogenesis to ensure increased phenotypic variability in offspring and improve chances of survival of the corresponding genetic lineage in conditions of changing environment ^51^. This hypothesis predicts that sperm DNA contains VMRs associated with genes involved in embryonic development and that VMRs non-specifically respond to stressors by an increase in methylation variation. This hypothesis also predicts that differential methylation in response to aging or stressors may result from increased variation in VMRs, although the mechanism of conversion of VMRs to DMRs remains unclear. To our knowledge, these predictions as well as the SEV hypothesis in general were never tested experimentally, and it is not clear if sperm epigenetic changes observed in many published studies may be explained by such a mechanism. Thus, the goal of the current study consists of (1) the testing of predictions from the SEV hypothesis using sperm DNA methylation data and (2) the development of evolutionary and molecular understanding of the nature of DNA methylation changes induced by aging and stressors.

## RESULTS

### VMRs are stable across ages and conserved in rats and mice

To identify VMRs in sperm of rodent models, we leveraged our previously published RRBS datasets ^8,52,53^ (Table 4). We identified 48,948 overlapping DNA methylation regions in 56- and 154-days-old mice. In this overlapping list, 2,566 and 3,248 regions were VMRs in 56- and 154-day-old mice, respectively, with 1,345 overlapping VMRs between age groups (Fig. 1A). In the rat dataset, 279,439 DNA methylation regions were identified in both 65- and 120-day-old animals. In this list, 9,872 and 17,418 regions were VMRs in younger and older rats, respectively, with 3,589 overlapping VMRs between age groups (Fig. 1B). In both species, VMRs identified in younger and older groups overlapped highly significantly with Fisher’s exact test p <0.00001. These overlaps were 8-fold and 6-fold higher in mice and rats, respectively, than overlaps predicted based on an assumption of independence of methylation variance in young and old animals. In both species, the amount of VMRs increased in older animals, suggesting age-induced gain in regions with high methylation variation. Notably, genes associated with VMRs overlapped between mice and rats (p <0.00001), indicating their conservation across species. Overall, these results suggest that concordant with the stochastic epigenetic variation hypothesis, DNA regions prone to methylation variation in sperm are not random and aging is a significant factor increasing the number of such regions.

**Fig. 1.**
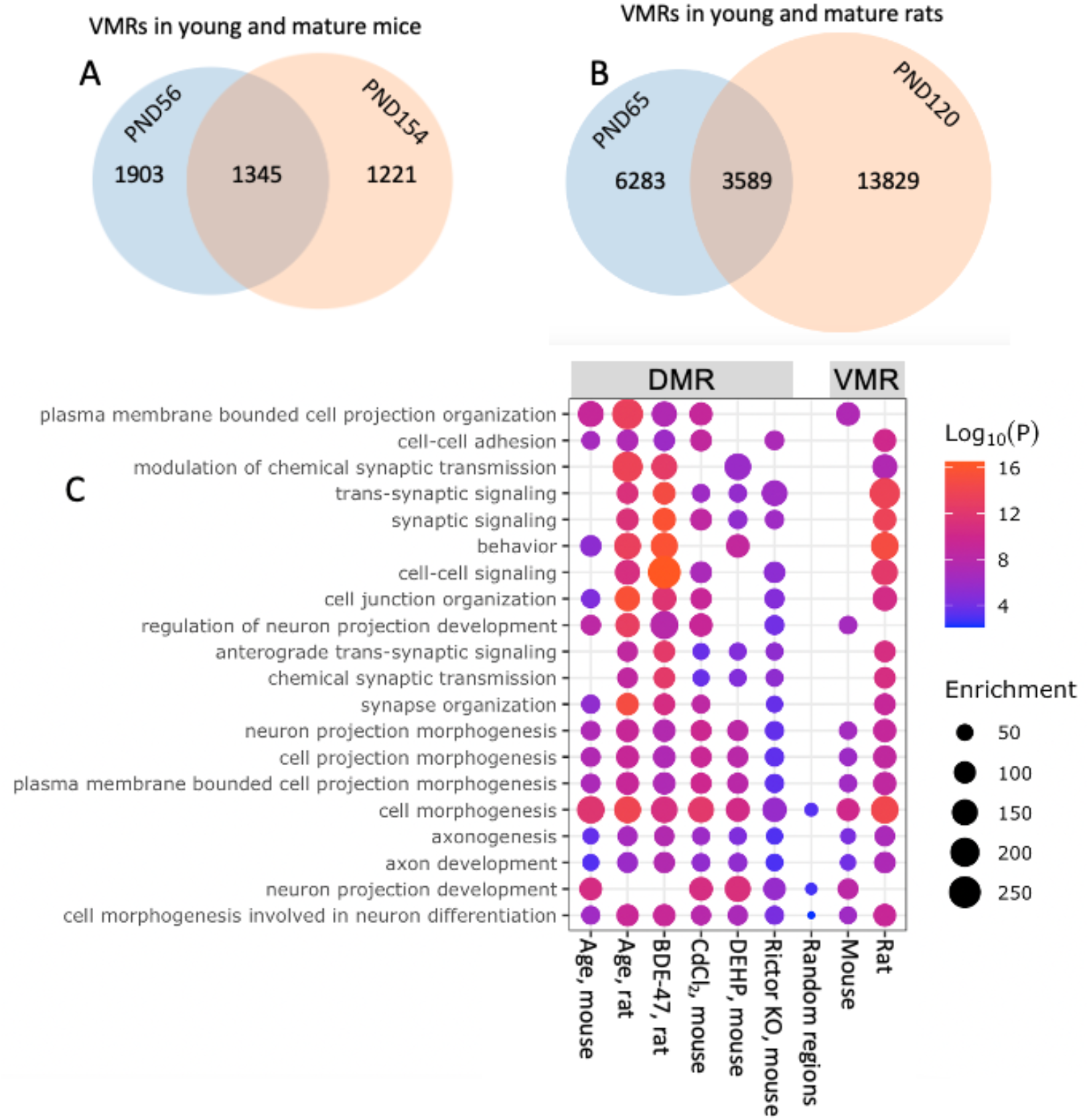
Sperm VMRs and DMRs in mice and rats. A-B, Overlap in sperm VMRs in animals of 2 ages in mice (A) and rats (B). C, Left to right, enrichment of biological categories in genes associated with sperm DMRs induced by aging in mice and rats, exposure to environmental pollutants BDE-47 in rats, cadmium in mice, and phthalates in mice, disruption of the BTB by Sertoli cell specific KO of Rictor in mice; 2,000 random DNA methylation regions; and VMRs in young mice and rats. Abbreviations: DMRs – differentially methylated regions, PND – postnatal day, VMRs – variable methylation regions.

### DMRs induced by age and stressors are enriched for developmental pathways

Similar sets of categories were enriched in DMR-associated genes in sperm in response to a broad range of factors (age, diet, chemical exposure, heat stress, psychological stress) in different species (human, rat, mouse) ^1,3,5–8,10,11,14,18–20,46–49^, suggesting the common mechanisms of DNA methylation response to various stressors. We analyzed enriched categories of genes associated with DMRs induced by age in mice and rats ^8,52,53^; environmental pollutants 2,2’,4,4’-tetrabromodiphenyl ether (BDE-47) (rats, perinatal exposure) ^8,52^, phthalates (mice, preconception exposure) ^20^, and cadmium chloride (mice, preconception exposure) ^11^; and blood-testis barrier (BTB) disruption by genetic manipulations in mice ^49^ (see Table 4 for the details of datasets used in this analysis). In all cases, DMR-associated genes were highly enriched with developmental categories including, for example, cell morphogenesis, neuron projection morphogenesis, synaptic signaling, axon development, and behavior (Fig. 1C). To test if RRBS approach selects for DNA methylation regions associated with developmental and neurodevelopmental genes, we conducted several enrichment analyses with 500 or 2,000 random DNA methylation regions identified by RRBS in mouse sperm. Random regions did not demonstrate enrichment for developmental and neurodevelopmental categories observed for DMRs responding to stressors and aging – see example in Fig. 1C. These later patterns suggest a possibility that the context of DNA regions determines the specific response of the sperm DNA methylation rather than the nature or timing of factors causing DNA methylation changes.

### VMRs are enriched for a similar set of developmental pathways

We used the lists of genes associated with significant VMRs in young mice and rats for enrichment analysis using Metascape. In both species, VMR-associated genes were highly enriched with developmental categories including, for example, cell morphogenesis, neuron projection morphogenesis, and axon development (Fig. 1C). Similar categories enriched in DMR- and VMR-associated genes in sperm suggest a functional link between DMRs and VMRs.

### Age-dependent DMRs overlap with VMRs

To test a hypothesis that age-dependent DMRs are a result of increased variation in VMRs, we analyzed the overlap between VMRs and DMRs in mouse and rat sperm using GSEA. First, age-induced methylation changes were identified across all DNA methylation regions. Next, VMRs in young animals were plotted against DNA methylation regions ranked by their percent of age-dependent methylation change. The results of these analyses are shown in Fig. 2A and B. Indeed, all significant VMRs grouped in the leftmost and rightmost regions of the differential methylation distribution, indicating that all DNA regions that undergo hypo- or hypermethylation with age correspond to VMRs in both species, while DNA regions not changing in methylation with age were depleted with VMRs.

**Fig. 2.**
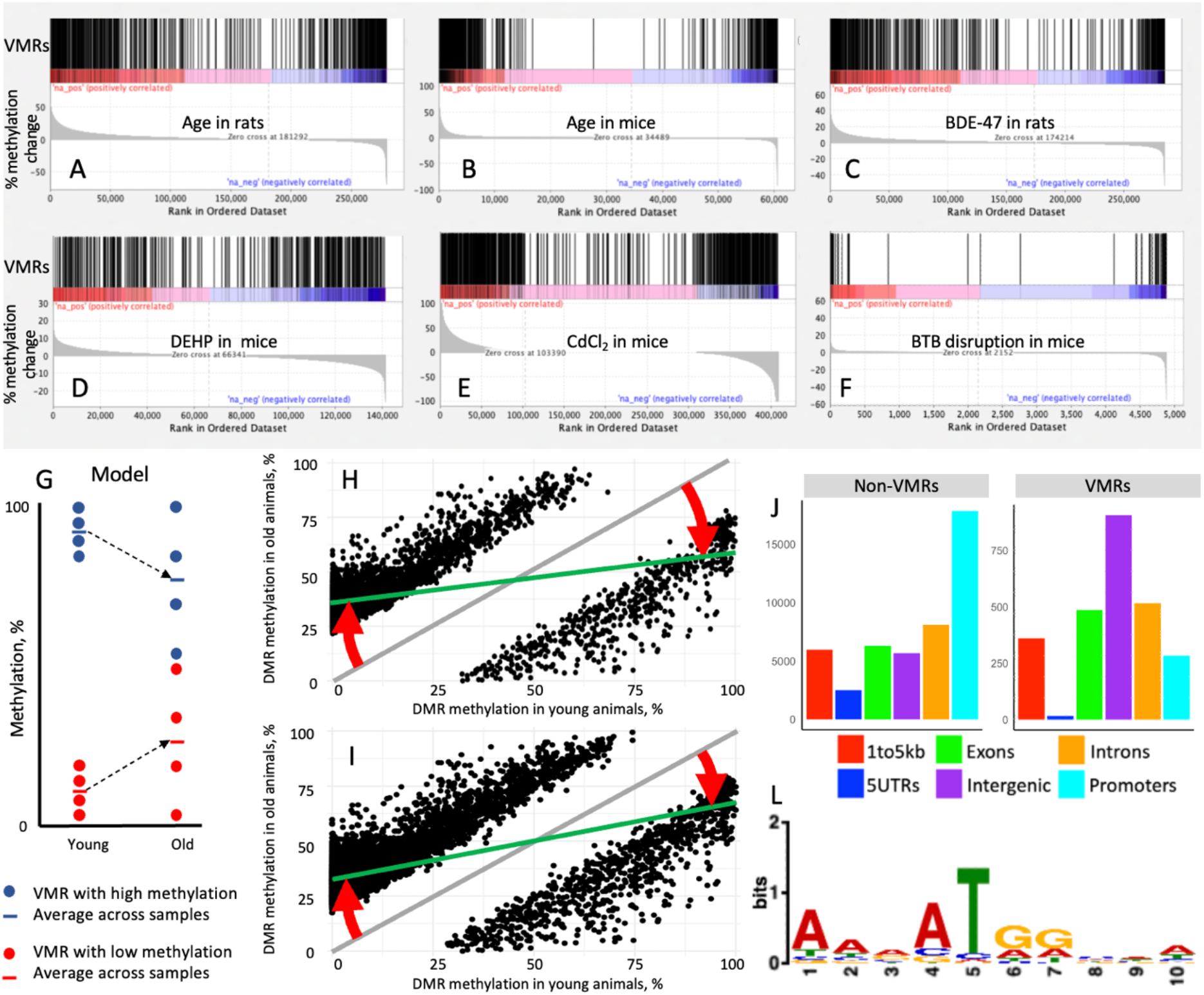
Characterization of VMRs. A-F, Position of rat (A, C) and mouse (B, D-F) VMRs in relation to DNA methylation regions ranked by methylation change induced by age in rats (A) and mice (B), exposure to BDE-47 in rats (C), exposure to DEHP (D) or cadmium (E) in mice and BTB disruption in mice (F). G, Mechanism that converts VMRs in DMRs: stress/aging induced stochastic changes in methylation of initially hypomethylated (red dots) and hypermethylated (blue dots) regions result in an asymmetric increase of variance and a shift in methylation values averaged for several (4 on this figure) organisms. H-J, Methylation values for age-dependent DMRs in young and mature mice (H) and rats (J) plotted against each other demonstrate that the dominant age-dependent change in methylation consists of increases in methylation of initially hypomethylated regions (red arrows in the left part of graphs) and decreases in methylation of initially hypermethylated regions (red arrows in the right part of graphs). In both graphs, each dot indicates one 100 bp age-dependent DMR; the grey diagonal line shows a hypothetical trendline for the case where methylation does not change with age; and the green line shows the real trendline for DMRs’ distribution. J, Overlap of sperm non-VMR and VMR regions with genomic elements in mice. L, Motif present in 90.5% of VMRs and significantly (p = 4.21e-04) enriched in VMRs as compared with non-VMRs in mouse sperm. Abbreviations: BDE-47 – 2,2’,4,4’-tetrabromodiphenyl ether, DEHP – Di(2-ethylhexyl) phthalate, DMRs – differentially methylated regions, VMRs – variable methylation regions.

### Stress-induced DMRs overlap with VMRs

We further hypothesized that increase in methylation variation in VMRs may also be a cause of differential methylation induced by stressors, such as chemical exposures and genetic perturbation affecting testis physiology. To test this hypothesis, we used the same GSEA approach as described in the previous paragraph. VMRs in young mice or rats were plotted against DNA methylation regions ranked by their percent methylation change induced by perinatal exposure to BDE-47 in rats, preconception exposure to phthalates or cadmium in mice, and BTB disruption via Sertoli-specific KO of Rictor in mice. Despite the very different nature and timing of these interventions, changes in DNA methylation occurred in VMR regions for all analyzed datasets (Fig. 2C-F). These results indicate that the genomic context of these regions determines non-specific response to various stressors, regardless of the type of stressor.

### DMRs are a result of the increased variability of VMRs

The observed link between VMRs and DMRs is unclear at first glance as VMRs respond to stressors by an increase of variation, while DMRs represent regions with increased or decreased methylation values, a seemingly different phenomenon. To understand the functional link between VMRs and DMRs, we developed the following hypothesis. Given that DNA methylation has a binary nature (presence or absence of methyl group on a CpG site), the majority of methylation values averaged for a DNA region, a population of cells, and/or organisms have values close to extreme methylation – namely approximately 0 or 100%. An increase in methylation variance in such regions will result in an asymmetric shift in averaged methylation values (Fig. 2G) as more cells will gain methylation in an initially hypomethylated site and more cells will lose methylation in an initially hypermethylated site. This model explains how an increase in the stochasticity of VMRs may result in DMRs. Indeed, the distribution of methylation values in age-dependent DMRs in young and mature mice and rats demonstrate that predominant age-dependent changes consist of an increase in methylation of sites hypomethylated in young animals and a decrease in methylation of sites hypermethylated in young animals (Fig. 2H and I).

### Overlap with repetitive elements does not characterize VMRs

Given that one of the most important functions of DNA methylation consists in suppression of repetitive elements, we analyzed if VMRs differ from other methylation regions in their association with these elements. We analyzed the overlap of VMRs and all methylation regions in DNA methylation datasets in young WT untreated mice and rats. Most repetitive elements were present in less than 1% of all DNA methylation regions and/or VMRs in both species. Thus, association with these elements cannot explain higher methylation variance in VMRs in comparison with other regions. Two types of repetitive elements were present in higher than 1% of regions in mice: low complexity DNA was found in 1.15% of all regions and 0.31% of VMRs, while simple repeats overlapped with 6.14% of all regions and 2.3% of VMRs. Thus, VMRs were 3.4-fold depleted with low complexity DNA and 2.7-fold for simple repeats with Fisher’s exact test p < 0.00001 for both element types. In rats, four types of repetitive elements were present in higher than 1% regions in all regions and/or VMRs: LINE were found in 1.21% of all regions and in 2.86% of VMRs, LTR were found in 2.56% of all regions and in 4.51% of VMRs, SINE were found in 2.96% of all regions and in 3.14% of VMRs, and simple repeats overlapped with 2.80% of all regions and 2.59% of VMRs. Out of these elements, VMRs were 2.35-fold enriched for LINE and 1.76-fold for LTR with Fisher’s exact test p < 0.00001 for both repetitive elements. The low percent of co-occurrence between methylation regions and repetitive elements in both VMR and non-VMR regions suggests that differences in methylation variance are likely not linked causatively to the presence or absence of repetitive elements.

### VMRs are depleted in open chromatin regions

We then hypothesized that higher variance in methylation observed in VMRs may be due to higher accessibility of corresponding regions for enzymes involved in methylation and demethylation of DNA. Given the DNA methylation profile of spermatozoa results from methylation/demethylation events that occur earlier in spermatogenesis, we tested if VMRs overlap with open DNA regions in spermatogonia, spermatocytes, spermatids, and mature sperm. For this analysis, ATAC-seq data for each cell type were extracted from ChIP-Atlas ^54–56^ and compared with 48,948 methylation regions in sperm, and enrichment/depletion of open chromatin regions were calculated for 2,566 VMRs as compared with 46,382 non-VMRs (Table 1). Unexpectedly, we observed that sperm VMRs are significantly (Fisher’s exact test p < 0.00001) depleted with open chromatin regions in all male germ cells. Interestingly, the number of VMRs associated with open chromatin is decreasing in the course of spermatogenesis. ATAC-seq is known to capture only hyper-accessible regions ^57^, thus depletion of VMRs in ATAC-seq regions suggests that their differential features may be due to binding with some biological molecules.

**Table 1.**
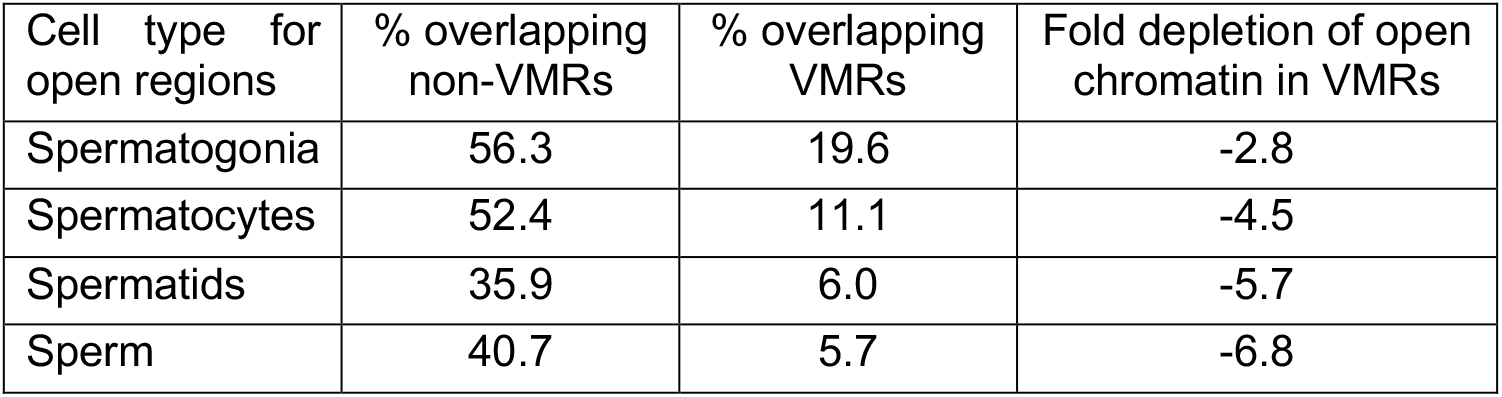
Overlap between open chromatin regions in spermatogenic cells and VMRs in mouse sperm.

| Cell type for open regions | % overlapping non-VMRs | % overlapping VMRs | Fold depletion of open chromatin in VMRs |
| --- | --- | --- | --- |
| Spermatogonia | 56.3 | 19.6 | -2.8 |
| Spermatocytes | 52.4 | 11.1 | -4.5 |
| Spermatids | 35.9 | 6.0 | -5.7 |
| Sperm | 40.7 | 5.7 | -6.8 |

### VMRs are differentially enriched with histones

Given that VMRs are depleted of open chromatin regions, we hypothesized next that VMRs may be associated with histones or specific modifications of histones. To test this assumption, we extracted ChIP-seq data for spermatogonia, spermatocytes, spermatids, and\ sperm from ChIP-Atlas ^54–56^. These datasets included data on the following histones: H2A.Z, H2AK119ub, H3, H3.3, H3K27ac, H3K27me3, H3K36me3, H3K4me1, H3K4me2, H3K4me3, H3K9ac, H3K9me2, H3K9me3, TH2A, and TH2B. DNA regions for these histones overlapped with 48,948 methylation regions in sperm and enrichment/depletion of histone-regions were calculated for 2,566 VMRs as compared with 46,382 non-VMRs for each germ-cell type (Table 2). We report only the data for these histone types which overlapped with >1% of either total methylation regions or VMRs or both in at least one cell type. These histone modifications include all histones, H3K27me3, H3K4me1, H3K4me2, and H3K4me3 (Table 2). Overall, these results demonstrate that VMRs are depleted of all histones throughout spermatogenesis. However, association of VMRs with specific modified histones is more complex.

**Table 2.** Overlap between DNA regions associated with histones in spermatogenic cells and VMRs in mouse sperm.

| Cell type for open regions | % overlapping non-VMRs | % overlapping VMRs | Fold depletion (-) / enrichment (+) in VMRs | Fisher's exact test p-value |
| --- | --- | --- | --- | --- |
| **All histones\*** |  |  |  |  |
| Spermatogonia | 38.8 | 28.3 | -1.35 | <0.00001 |
| Spermatocytes | 41.7 | 40.9 | -1.02 | 0.41 |
| Spermatids | 36.6 | 28.5 | -1.28 | <0.00001 |
| Sperm | 38.9 | 18.6 | -2.04 | <0.00001 |
| **H3K27ac** |  |  |  |  |
| Spermatogonia | 3.4 | 2.5 | -1.31 | 0.02 |
| Spermatocytes | 0.83 | 0.23 | -3.42 | 0.0002 |
| Spermatids | 0.74 | 0.12 | -6.05 | <0.00001 |
| Sperm | 0.06 | 0 | -- | 0.40 |
| **H3K27me3** |  |  |  |  |
| Spermatogonia | 2.2 | 4.2 | 1.85 | <0.00001 |
| Spermatocytes | 5.7 | 2.9 | -1.91 | <0.00001 |
| Spermatids | 6.6 | 11.8 | 1.73 | <0.00001 |
| Sperm | 5.3 | 3.2 | -1.62 | <0.00001 |
| **H3K4me1** |  |  |  |  |
| Spermatogonia | 0.11 | 0.23 | 1.91 | 0.13 |
| Spermatocytes | 2.4 | 5.7 | 2.25 | <0.00001 |
| Spermatids | 5.6 | 5.7 | 1.01 | 0.93 |
| Sperm | 0.02 | 0 | -- | 1 |
| **H3K4me2** |  |  |  |  |
| Spermatogonia | 2.5 | 6.5 | 2.40 | <0.00001 |
| Spermatocytes | 2.1 | 2.0 | -1.04 | 0.89 |
| Spermatids | 2.9 | 1.9 | -1.53 | 0.001 |
| Sperm | 17.9 | 9.6 | -1.82 | <0.00001 |
| **H3K4me3** |  |  |  |  |
| Spermatogonia | 30.7 | 14.7 | -2.04 | <0.00001 |
| Spermatocytes | 30.1 | 29.1 | -1.03 | 0.31 |
| Spermatids | 20.7 | 8.7 | -2.31 | <0.00001 |
| Sperm | 14.8 | 4.6 | -3.10 | <0.00001 |

\* Sum of H2A.Z, H2AK119ub, H3, H3.3, H3K27ac, H3K27me3, H3K36me3, H3K4me1, H3K4me2, H3K4me3, H3K9ac, H3K9me2, H3K9me3, TH2A, and TH2B.

Given that the most significant DNA methylation changes in the course of spermatogenesis occur at spermatocyte ^23–25^ and spermatid stages ^26^, association of VMRs with histones during these stages may be more important. In spermatocytes, VMR regions are significantly depleted with H3K27me3 and H3K27ac and significantly enriched with H3K4me1. Spermatid VMRs were depleted of H3K27ac and H3K4me2/3 and enriched with H3K27me3.

H3K4me1 is a well-recognized mark of poised or active enhancers while H3K27ac is a mark of active enhancers only ^58,59^. H3K27me3 is a major repressive histone modification usually attributed to facultative heterochromatin ^60^. Finally, H3K4me2 and H3K4me3 mark chromatin accessible for transcription and are usually associated with active 5′ end of genes and promoters, respectively ^61^. These results demonstrate that VMRs are associated with distinctly different profiles of modified histones than non-VMRs. However, given only a small fraction of VMRs is associated with differentially enriched/depleted histone variants, it is unlikely that these associations causally determine the variable nature of VMRs.

### VMRs distribution across genomic elements and motifs

To analyze if VMRs are differentially associated with different genomic features, mouse VMRs and non-VMRs were annotated to the genomic features of the mouse genome. The results of this analysis are shown in Figure 2J. Non-VMR DNA methylation regions captured by RRBS are enriched in promoters, while the highest number of VMRs’ overlap is with intergenic regions.

To test if VMRs are characterized by specific binding sites, we submitted the lists of VMR and non-VMR sequences in the sperm of young untreated mice to the STREME tool ^62^. This analysis identified 14 differentially enriched motifs with Fisher Exact Test p < 0.05 (Table 3). Only 5 of these motifs had an *E*-value (Fisher Exact Test *p*-value adjusted for multiple comparisons) < 0.05. Given that the goal of this analysis is to identify discriminative features of VMRs, we focused further on the only significant (E-value < 0.05) motif (AARATGGMWW) that was present in the majority (90.5%) of VMRs. Subsequently, we submitted the selected motif to the Tomtom tool of the MEME Suite and identified 21 matching motifs in the HOCOMOCO Mouse (v11 CORE) database, corresponding to binding sites of the following proteins: ZFP42, NR4A1, MEF2A, MEF2C, MEF2D, TAF1, YY1, PRDM9, SRF, FOXL2, OTX2, ATOH1, IRF3, NKX6-1, NEUROD2, NEUROD1, ISL1, ALX1, SOX10, ELF5, MSGN1, listed by descending significance. Binding sites of these proteins matched our VMR motif (AARATGGMWW, Fig. 2L) with p<0.05, however, both q-value and E-value calculated to adjust for multiple comparisons were >0.05 for every protein. ZFP42/REX1 binding site had the highest significance of match (p = 4.21e-04) to the motif enriched in the highest number (90.5%) of VMRs.

**Table 3.**
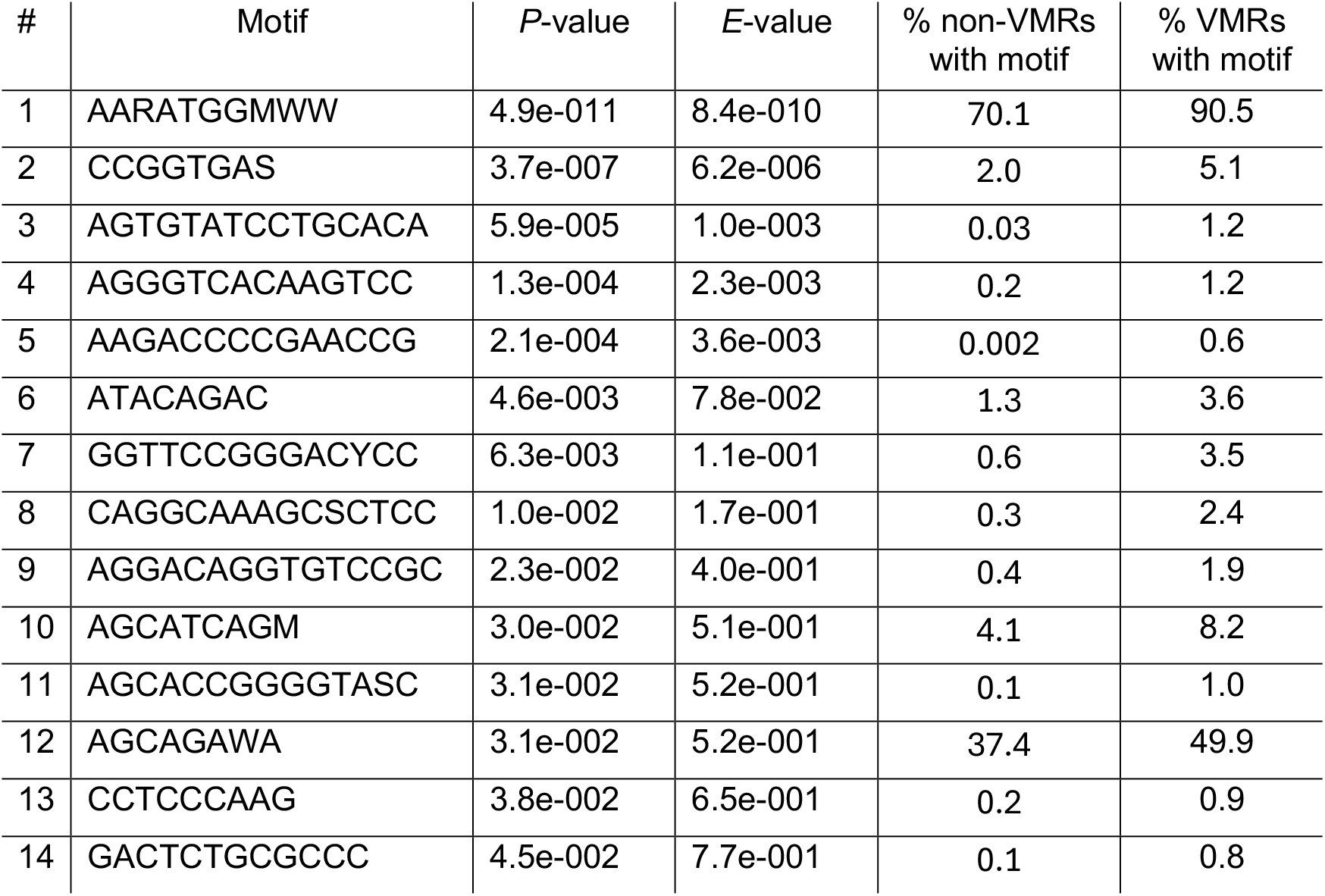
DNA motifs enriched in mouse sperm VMRs.

**Table 4.**
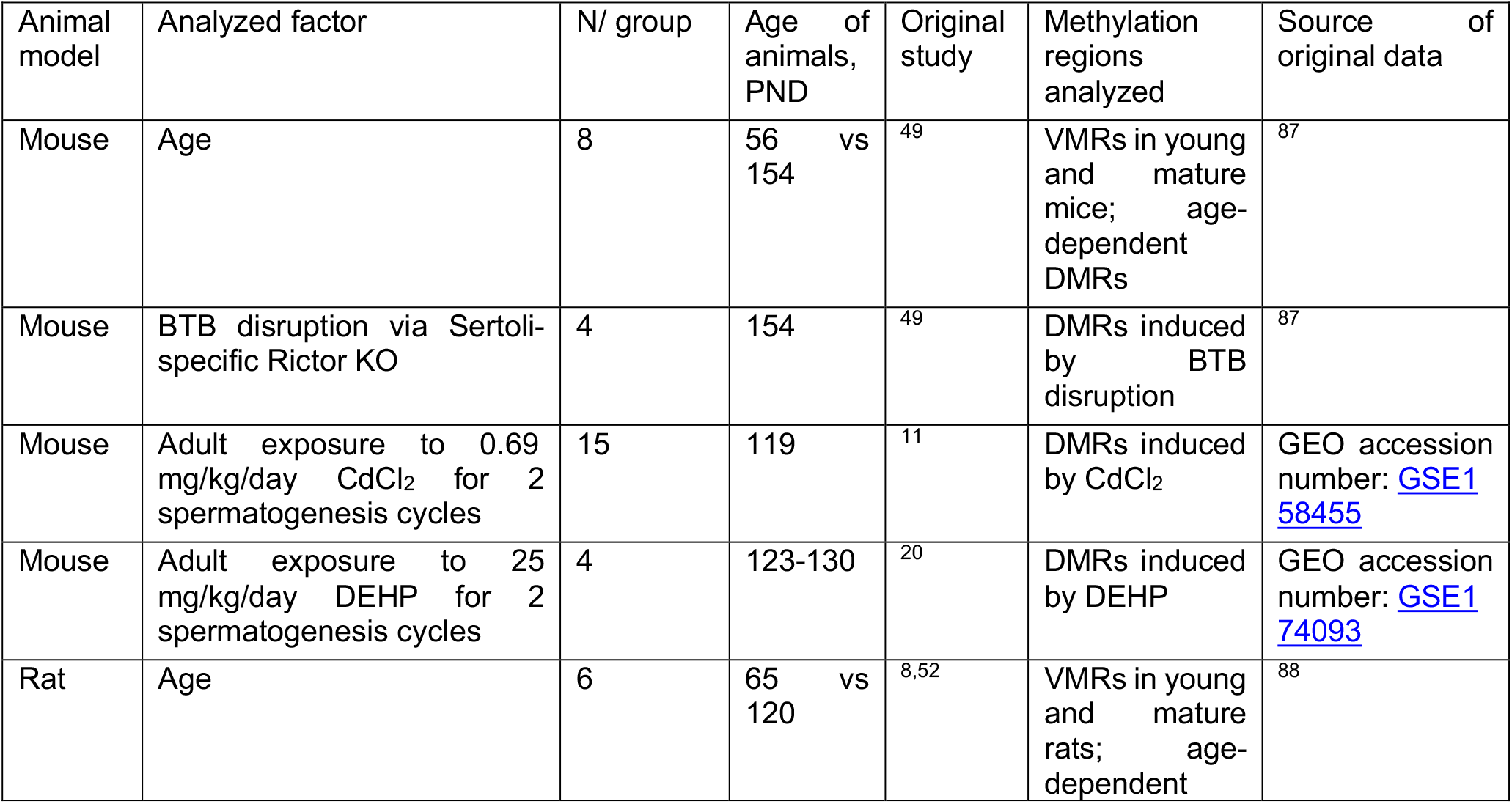
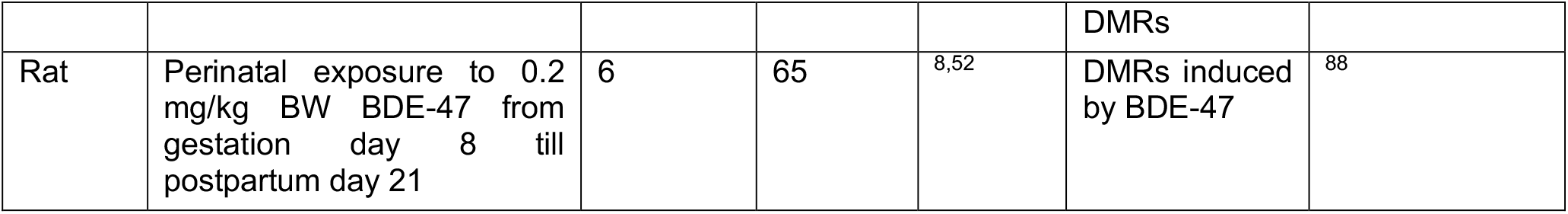
Sperm RRBS datasets used in this study to identify VMRs and DMRs.

ZFP42/REX1 is recognized as an epigenetic remodeler and a marker of pluripotency ^63^. Another epigenetic remodeler binding the same motif is the histone methyltransferase, PRDM9. Proteins of other matching motifs are transcription factors essential for muscle development (MEF2A, MEF2C, MEF2D), nervous system development (NEUROD1, NEUROD2, SOX10, OTX2, ATOH1), mesoderm differentiation (MSGN1), keratinocyte differentiation (ELF5), development of craniofacial structures (ALX1), cell morphogenesis (SRF), cell cycle regulation\ (Taf1), gonad differentiation (FOXL2), type I interferon response (IRF3), development of insulin-producing beta cells and insulin secretion (NKX6-1, ISL1), and stress response cell differentiation, apoptosis, proliferation, inflammation, and metabolism (NR4A1)^64^.

### Overlap of VMRs with regions escaping methylation reprograming

Given that VMRs are enriched with ZFP42/REX1 binding motifs and that ZFP42/REX1 plays a role in imprinting, we hypothesized that VMRs may represent regions related to imprinting phenomena. To test this hypothesis, we first compared the overlap of genomic coordinates of VMRs with coordinates of imprinted genes in mice and rats. The lists of imprinted genes were taken from the Geneimprint database ^65^ and included 159 mouse and 55 rat genes. No overlap with sperm VMRs in either species was identified. Next, we compared the coordinates of mouse sperm VMRs with regions that escape epigenetic reprogramming during early embryogenesis. Regions that escape demethylation were identified as regions with >15% methylation in blastocyst and regions that escape remethylation were identified as regions with <15% methylation in both spermatozoa and embryonic stem cells. Our analysis demonstrated that VMRs are not enriched with either type of escapees.

## DISCUSSION

This study identified sperm VMRs as the material basis for DNA methylation changes in response to a broad range of stressors and aging. We demonstrate that the genomic context of DNA regions determines the response of sperm methylome to various factors rather than the type of stressor and/or timing of these factors. Because methylation variation in VMRs increases with age, sperm methylome response to stressful factors may be characterized as the acceleration of epigenetic aging.

VMRs in mice and rats are conserved and are associated with genes enriched for developmental categories. These characteristics match predictions from the stochastic epigenetic variation hypothesis, suggesting that VMRs represent an evolutionary adaptation that increases the chances of survival of the genetic lineage in conditions of changing environment ^51^. VMRs may represent stochastic switches in branching points of developmental trajectories of Waddington’s epigenetic landscape ^66^. That view of epigenetic changes in sperm suggests that phenotypic changes in offspring conceived by older fathers or fathers exposed to stressful environments are characterized by increased phenotypic variation, rather than by directional shift in phenotypic traits. Experimental testing of this hypothesis is needed in future research.

Age-dependent changes in sperm DNA methylation may have a similar nature in somatic cells and may explain mechanisms behind epigenetic clocks. Indeed, it was demonstrated that many cytosines increase in methylation variation in human somatic cells with age ^67–71^ and VMRs were identified in human whole blood samples ^72^ and mouse and human livers ^51^, mouse blood ^73^, and human brain ^51^. It was also shown that human blood VMRs are enriched in enhancer regions ^68^. Recent single-cell methylation analysis using mouse stem cells demonstrated that major age-dependent changes in DNA methylation are characterized by a stochastic increase in methylation of initially hypomethylated regions and a decrease in hypermethylated regions ^73^. It was also shown using purely simulated data that accumulation of stochastic variation is sufficient to build epigenetic clocks that capture age-dependent shifts in DNA methylation ^45^. Additionally, it was shown that changes in bulk blood DNA methylation throughout functional aging (3-35 months of age) in mice are small, with the largest change around 30% per CpG ^73^. This observation is concordant with our model (Fig. 2G), suggesting that age or stress-dependent changes in DNA methylation are a result of increasing asymmetric variation across a population of cells, rather than the result of directional and coordinated change. Indeed, if the initially hypomethylated region undergoes a coordinated directional shift in methylation, the maximum shift in methylation of approximately 100% may be achieved when all cells gain methylation. In the case of methylation shift due to increasing variation, the maximum methylation shift may only reach 50% - the point when variance is maxed out (max entropy, min information).

We tried to elucidate if VMRs may be involved in the intergenerational transfer of epigenetic information as suggested by the stochastic epigenetic variation hypothesis. We observed that mouse sperm VMRs are enriched with ZFP42/REX1 motif (Table 3). This gene is highly expressed only during spermatogenesis (in spermatogonia and spermatocytes) and in early embryo, specifically in placenta ^63^ - extra-embryonic tissue mostly controlled by paternally expressed imprinted genes ^74^. Although ZFP42/REX1 is not expressed in the late embryo, its gene dosage during early embryogenesis is critical for the survival of the late-stage embryos and neonates, suggesting that during spermatogenesis ZFP42/REX1 creates some epigenetic signature that instructs embryo development at later stages ^63^. This assumption is supported by the fact that paternal inheritance of deactivated ZFP42/REX1 causes higher levels of lethality in embryos than maternal inheritance ^63^. It is also the central negative regulator of X chromosome inactivation during early female embryogenesis ^75^ and a central positive regulator of X chromosome reactivation ^76^. ZFP42/REX1 also plays a role in genomic imprinting and is hypothesized that it binds unmethylated allele of imprinted genes and protects them from DNA methylation removal during embryogenesis ^63^.

The association of VMRs with ZFP42/REX1 motif suggests that these regions may survive methylation reprogramming in the early embryo and affect developmental trajectories. However, we did not find any significant overlap between VMRs and methylation reprograming escapees. Thus, it remains unclear if DNA methylation in sperm VMRs has a higher propensity to contribute to the developmental trajectories in offspring than methylation in non-VMR regions.

It is also important to mention here that the ability of sperm methylation patterns to affect initial stages of embryo development before being reprogramed during blastocyst is poorly understood. According to the classical view, the first transcription in newly formed embryos starts in the eight-cell stage, around 3 days after fertilization ^77^. Recent research demonstrated, however, that embryonic transcription initiates as early as at the one-cell stage in humans ^78^ and mice ^79^, and that zygotic transcription activation starts from the paternal genome ^80^. These reports taken together suggests that paternal epigenome may play an important role in determining the developmental trajectories of the embryo before the global wave of methylome reprogramming.

We also attempted to identify specific features of VMRs that may be causatively linked to their high methylation variation. Although VMRs were differentially enriched with some repetitive elements and histone modifications, the fact that both modified histones and repetitive elements overlap with only a small portion of VMRs suggests that these features cannot explain causatively the stochastic nature of DNA methylation in these regions. Surprisingly, VMRs were depleted with open chromatin regions in male germ cells, indicating that these regions are bound to some biological molecules (Table 1). However, they were mostly depleted with histones as well (Table 2), suggesting that molecules other than histones protect VMRs from hyperactive transposase that is the central element of ATAC-seq technology. We hypothesize that association with ZFP42/REX1 may play a role in the stochastic instability of methylation in VMRs. Indeed ZFP42/REX1 gene has been duplicated from YY1 in the Eutherian lineage ^81^ and both transcription factors still share similar binding motifs. The two proteins competing for the same binding site may have different effects on DNA methylation of corresponding regions. For example, ZFP42/REX1 was shown to protect from methylation the paternal allele of imprinted *Peg3* gene ^63^, while YY1 promotes methylation of *Peg3* during oogenesis ^82^. Additionally, ZFP42/REX1 itself may have a dual effect on DNA methylation of regions to which it is bound. On one hand, it may shield them from methylation; on another hand, some evidence indicates that ZFP42/REX1 can recruit *de novo* DNA-methyltransferase DNMT3b ^83^ and increase *Dnmt1* expression ^63^. Thus, these two mechanisms – competition between ZFP42/REX1 and YY1 for the same binding sites and their opposite effects on methylation, as well as the potentially dual role of ZFP42/REX1 itself – may be responsible for high methylation variation in VMRs. This hypothesis is to be tested experimentally in future research.

## Conclusions

This is the first study to identify sperm VMRs as the material basis for DNA methylation changes in response to a broad range of stressors and aging. These findings are concordant with the stochastic epigenetic variation hypothesis, which proposes that VMRs result from adaptive evolution by increasing the chances of genetic lineage survival in changing environmental conditions. Our data suggest that high level of stochastic variation in DNA methylation within VMRs may be causally linked to ZFP42/REX1 protein, an epigenetic remodeler involved in genomic imprinting. The nature of stochastic instability of VMRs, as well as their ability to deliver epigenetic information to offspring, requires additional research.

## MATERIAL AND METHODS

### Datasets

This study is based on sperm DNA methylation datasets previously published by our and other groups (Table 4). To achieve uniformity of data analysis we used only studies using reduced representation bisulfite sequencing (RRBS) as the major method for DNA methylation analysis and reanalyzed all data using a pipeline described below. The following datasets were used to analyze changes in sperm DNA methylation in response to aging and other stressful factors. To identify VMRs in young untreated wildtype mice we used RRBS data from 8 individual 56-day-old animals from our recent study ^49^. DNA methylation profiles of 56 and 154-day-old mice from the same study ^49^ were used to identify age-dependent changes in mouse sperm epigenome. The original study ^49^ demonstrated that the absolute majority of DNA methylation regions undergo semi linear increase or decrease in DNA methylation during the studied period from postnatal day 56 till 334. Thus, a comparison of any two age groups within this period may be used to characterize sperm methylome aging. Additionally, this study was the source of RRBS datasets representing wildtype (WT) mice and mice with Sertoli-specific knockout of Rictor – a genetic manipulation resulting in a disruption of the blood-testis barrier (BTB) ^49,84^. To analyze age-dependent changes in sperm methylome in rats we used our RRBS data from 65 and 120-day-old animals from previously published research ^8,52^. The same dataset was used to analyze the effects of perinatal exposure to 0.2 mg/kg/day of a brominated flame retardant 2,2’,4,4’-tetrabromodiphenyl ether (BDE-47) which was conducted via pipette feeding of pregnant/lactating dams to a solution of BDE-47 in corn oil or vehicle staring from gestational day 8 till post-partum day 21. Changes in mouse sperm methylome in response to phthalate exposure were analyzed using data from our previous study ^20^. In this study, sperm was obtained from 17-18 weeks old animals following continuous exposure to 25 mg/kg/day di(2-ethylhexyl) phthalate (DEHP), a compound found in plastics and personal care products, or vehicle control (50% dimethyl sulfoxide and 50% ethanol) via subcutaneous osmotic pumps for the duration of 2 spermatogenesis cycles. Changes in mouse sperm methylome in response to cadmium exposure were analyzed using data from a published study ^11^ in which animals were exposed to approximately 0.69 mg of CdCl_2_ per kg of body weight for 9 weeks via drinking water. Data on mouse sperm ATAC-seq and CHIP-seq with antibodies for various histones were taken from ChIP-Atlas a data-mining suite powered by full integration of public sequencing data ^54–56^. RepeatMasker ^85^ was used to identify DNA sequences for interspersed repeats and low-complexity DNA sequences. The lists of imprinted genes were taken from the Geneimprint database (https://www.geneimprint.com) ^65^. Finally, the list of genomic regions that survive epigenetic reprogramming in mice was taken from the study which provides a comparison of DNA methylation values in sperm, oocytes, blastocyst, and embryonic stem cells ^86^.

### Identification of DMRs

Raw RRBS reads were trimmed using TrimGalore (v 0.6.6) and a NuGEN-specific adaptor trimming scripts available from GitHub (nugentechnologies/NuMetRRBS). Trimmed reads were mapped using Bismark-Bowtie2 with no mismatch allowed. GRCm38/mm10 and mRatBN7.2/rn6 genome assemblies were used to map mouse and rat data respectively. Methylation counts were called using Bismark extract. Differentially methylated regions were identified using the Methyl kit (v 1.24.0) pipeline ^89^. In brief, the genome was tiled into sliding windows and a weighted methylation level was calculated for each window. To minimize error in base calling, we filtered out bases with less than 10x coverage and read counts more than 99.9^th^ percentile, and coverage values were normalized using default settings. We used a logistic regression model for *p*-value calculation subsequently adjusted for multiple comparisons (FDR) using the SLIM method for final DMR identification. Individual DMRs were identified for a 100 bp sliding window with q < 0.05.

### Identification of VMRs

Methylation variance was calculated for each 100 bp region for mouse (56 and 154 days of age, n = 8 per age) and rat (65 and 120 days of age, n = 6 per age) sperm DNA (Table 4). Top VMRs were then identified for each age/species as regions with the highest values of interindividual variance, using a method for the identification of cut-off points in descriptive high-throughput omics studies ^90^. This approach assumes that the variance of DNA regions follows generally biphasic distribution with the majority of regions having low variance and a smaller number of regions having high variance. The method identified the threshold between low-variance and high-variance as a maximum inflection point in a ranked distribution of all regions.

### Enrichment analyses

Fisher’s exact test was used to test the significance of overlaps of different genomic regions (VMRs, DMRs, ATAC-seq, ChIP-seq, repetitive elements, regions escaping methylation reprogramming, and genes associated with VMRs in different species). RRBS method used for DNA methylation analysis may introduce some biases as it selects only for regions enriched with CCGG sequence. To avoid potential biases introduced by RRBS, we conducted all enrichment analyses only for regions that were identified by RRBS. To identify fold enrichment/depletion of VMRs with other genomic regions (VMRs, DMRs, ATAC-seq, ChIP-seq, repetitive elements, regions escaping methylation reprogramming, and genes associated with VMRs in different species) we compared the observed number of overlapping elements with the calculated number of overlapping elements based on the probability of overlap if no interaction between the two elements exists (null hypothesis).

Additionally, gene set enrichment analysis (GSEA) was used to visualize overlaps between DMRs and VMRs and establish if DMRs are enriched with VMRs ^91^. For functional enrichment analysis DMRs or VMRs were assigned to the closest gene (<5 kb upstream transcription start site, promoter, 5’UTR, exon, or intron) or intergenic region. Spatial genomic annotation was conducted using annotatr package (v1.24.0) ^92^ and annotated to the genomic features of the Ensembl genome. Genomic features were compared using the GenomicRanges package (v1.50.2) ^93^. Metascape ^94^ analysis was used to analyze biological categories associated with genes annotated to DMR or VMRs.

### Motif analysis

The lists of VMR and non-VMR (control) regions were submitted to STREME tool ^95^ of the MEME Suite 5.5.5 ^62^ to identify DNA motifs significantly overrepresented in VMRs. These motifs were submitted to the Tomtom tool of the MEME Suite to identify matching annotated motifs in the HOCOMOCO Mouse (v11 CORE) database. Given that DNA motifs are not yet well annotated in the rat genome, this analysis was done for mice only.

